# ANOMALY: A Snakemake pipeline for identifying NuMTs from Long-Read Sequencing Data

**DOI:** 10.1101/2025.04.08.647704

**Authors:** Nirmal Singh Mahar, Rachit Singh, Ishaan Gupta, Shweta Ramdas

## Abstract

**Motivation:** Nuclear mitochondrial DNA segments (NuMT) can significantly affect cellular processes, including cancer development and disease progression. Current methods to call NuMTs rely on short-read sequencing data but struggle to resolve complex NuMTs. These limitations can be overcome by employing long-read sequencing data. However, no such workflow exists to capture NuMTs from long-read sequencing data.

**Results:** Here, we introduce ANOMALY, a novel, easy-to-use workflow for detecting NuMTs from long-read sequencing data. The pipeline takes raw sequencing data or aligned data and calls and visualizes sample NuMTs. On 50 simulated datasets, the pipeline demonstrated high accuracy, with a precision of 1.000, a recall of 0.989, and an F1-score of 0.994. The pipeline underscores the limitations of short-read data in resolving and capturing complex NuMTs while demonstrating that long-read data enables their accurate identification.

**Availability and Implementation:** The Snakemake pipeline employs Python, Bash and R and is published under an open-source GNU GPL v3 license. Detailed information about setting up and running the pipeline and the source code can be accessed at https://github.com/Nirmal2310/ANOMALY.

## 1. Introduction

Nuclear mitochondrial DNA segments (NuMTs) represent segments of mitochondria-derived DNA in the nuclear genome. These insertions are estimated to occur at a rate of ~ 5 × 10^−6^ per base pair per generation in humans^1^. A recent study utilising whole genome sequences from 66,083 samples has determined that each individual carries 4.7 NuMT events on average^2^. These NuMTs have been implicated in genome function^3^ and disease^4–7^, and used in phylogenetic inference^8^.

Existing tools for calling NuMTs from whole genome sequencing data, including DINUMT^9^ and the approach proposed by Wei et al.^10^, are based on short paired-end reads only. With the increasing adoption and availability of long-read sequencing technologies in both small and population-scale genomic studies, there is a need for a tool that can characterise NuMTs from these datasets. Long reads offer higher accuracy to call structural variants^11^, including long insertions like NuMTs; several tools have been built specifically for this purpose and have led to the discovery of novel structural variations in human populations. However, no such tool exists specifically for detecting NuMTs alone, and such tools alone are insufficient to accurately call NuMTs without many additional steps, each of which requires parameter tuning.

We present a novel Snakemake-based pipeline called ANOMALY (**<u>A</u>**nalysis of **<u>N</u>**uclear inserts **<u>O</u>**f **<u>M</u>**itochondri**<u>A</u>** using **<u>L</u>**ong-read sequencing in **<u>Y</u>**our data) for NuMT calling from long-read whole-genome sequencing data. ANOMALY is a pipeline based on existing state-of-the-art open-source methods. Implemented in Snakemake^12^, the tool (with its necessary dependencies) is easy to install and quick to run. In simulated data based on the T2T-CHM13 genome^13^, ANOMALY identified 536 out of 542 simulated NuMTs, with a false negative rate of 0.011 (Table 1, Supplementary Table 1).

## 2. Method

### Overview

ANOMALY is a UNIX-based Snakemake pipeline that takes as input raw sequencing data, or whole-genome sequenced aligned data (Figure 1A). Using the existing structural variant (SV) caller Sniffles2^14^, the pipeline first calls all structural variants (SV) in the sample (step 1). The sequence of each identified insertion is then aligned to the concatenated reference mitochondrial genome FASTA using Nuclear BLAST^15^. The concatenated reference mitochondrial FASTA file used for this local alignment is generated using SeqKit^16^ to account for the circular mitochondrial genome and capture the NuMTs comprising the mitochondrial genome’s control region. Potential NuMTs identified by Nuclear BLAST are then filtered and prioritised based on the E-Value (<1e-3) and query coverage (>=70%). All of the above Sniffles2 and Nuclear BLAST parameters were optimised using simulated data to minimise false positive rates. However, users can change these parameters by editing the config file provided with the tool. The final output of ANOMALY contains information about the nuclear breakpoint of the NuMT, including the size and location of the mitochondrial genome part integrated into the nuclear genome and a circos plot showing a visual representation of all integrated NuMTs (Figure 1B).

**Figure 1.**
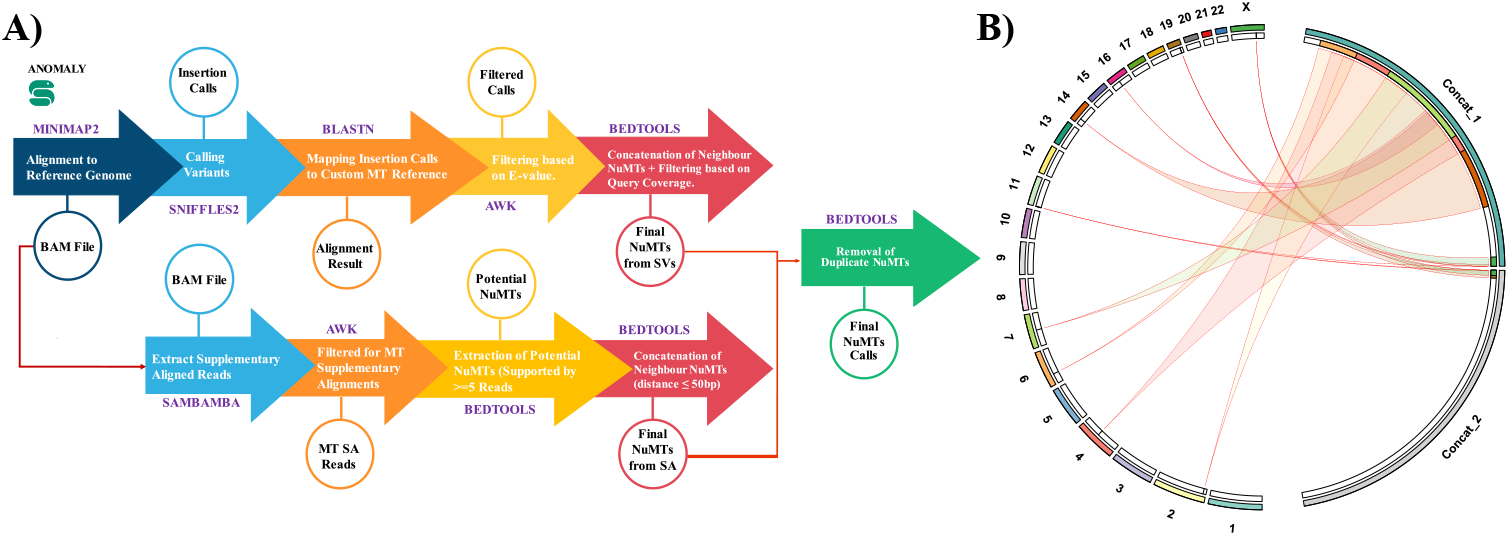
**A)** ANOMALY workflow. The workflow is implemented as a Snakemake pipeline designed to detect NuMTs. It accepts raw sequencing data in FASTQ format or pre-aligned data in BAM format as input. The pipeline produces a TSV file containing NuMT calls and visual representations as a Circos plot, saved in PNG and SVG formats. In the schematic representation, the open-source tools used in the pipeline are highlighted in purple. The steps are described along the arrows connecting the workflow components. Outputs of each step are indicated within circles. **B)** The Circos plot illustrates the integration of Nuclear Mitochondrial DNA Sequences (NuMTs) across various chromosomes in the nuclear genome. The outermost circular segments represent individual chromosomes (1–22 and X) and concatenated mitochondrial genome segments (Concat_1 and Concat_2). The curved links in the inner region indicate the insertion events of NuMTs from the mitochondrial genome into nuclear chromosomes. To visualise the NuMTs containing the control region of the mitochondrial genome, we have utilised the concatenated genome of Mitochondria (shown as Concat_1 and Concat_2). Hence, all the NuMTs containing control regions will be split into two links, one coming from the end of the first mitochondrial segment (Concat_1) and the second coming from the start of the second mitochondrial segment (Concat_2).

### 2.1: Alignment with the Reference Genome

If the user chooses to input unaligned FASTQ files to the tool, the pipeline first aligns the input data to the human reference genome (including reference mitochondrial genome) using Minimap2 version 2.28 with parameters optimised for long reads (allowing soft clipping for supplementary alignments), followed by sorting and indexing of aligned data using SAMtools version 1.17^17^.

### 2.2 Identification of Potential NuMTs from Structural Variants

The sorted and indexed BAM file is then used to call Structural Variants (SVs) using Sniffles version 2.5 with parameters “sniffles --minsupport 4 --mapq 1 --minsvlen 15 --genotype-ploidy 2”. The insertion calls identified by Sniffles are then extracted and realigned with the concatenated reference mitochondrial FASTA using BLASTn version 2.15.0. The concatenated FASTA is generated using the concat command of SeqKit version 2.9.0. The main purpose of using the concatenated FASTA for the realignment is to capture NuMTs covering the control region of the mitochondrial genome. All the potential NuMTs captured by the realignment are then filtered based on E-value (<1e-3). The filtered NuMTs are concatenated if the distance between the calls is less than 10 bp using BEDtools version 2.30.0^18^. The concatenated NuMTs are filtered based on query coverage (>=70%), resulting in the final NuMT calls from Structural Variants.

### 2.3 Identification of Potential NuMTs from Supplementary Alignments

In the previous study, it was reported that the size of the NuMTs can range from 39 bp to the complete length of the mitochondrial genome^9^. Larger NuMTs are difficult to capture as Structural Variants as this often requires ultra-long reads as an input, which results in additional cost^19^. However, by leveraging the information stored as Supplementary alignments in the BAM file, this tool can capture full-length NuMTs. The pipeline first extracts the aligned reads from the nuclear genome having supplementary alignments with the mitochondrial reference genome. The neighbouring regions in the nuclear genome are concatenated using BEDtools if the distance between them is less than 50 bp. The potential nuclear breakpoints are then determined using the number of supporting reads (>=5). The final NuMT calls from supplementary alignments are then reported based on the breakpoints supported by at least 5 reads.

### 2.4 Final NuMT Call and Visualisation

The NuMT calls from structural variants and supplementary alignments are concatenated, and unique calls are retained to report the final NuMTs. The final NuMT calls are then visualised as a circos plot using the R package circlize^20^ version 0.4.16. The resulting circos plot is saved locally in PNG and SVG format. At the end of the run, the pipeline will produce one single circos plot annotating all the NuMTs called in all the input samples to give a more comprehensive overview of the NuMTs.

#### Data Simulation

To evaluate the pipeline’s performance, we simulated 542 unique NuMTs across 50 samples, with NuMT size ranging from 30 bp to the full human mitochondrial genome length (16,569 bp). The CHM13 human reference genome was modified using the ‘mutate’ command in SeqKit version 2.9.0 to introduce the defined insertion sequences. Subsequently, PBSIM3 version 3.0.4^21^ was employed to generate sequencing data from the modified CHM13 genome using the ERRHMM-RSII error model. The resulting dataset had a 30X coverage depth and an average read length of 10 Kbp. The pipeline was then used to detect NuMTs in each simulated sample, and the identified NuMTs were compared to the known true calls to establish true positives (TP), false positives (FP) and false negatives (FN). Finally, precision, recall and F1-score were calculated from these metrics to assess the overall performance of the pipeline.

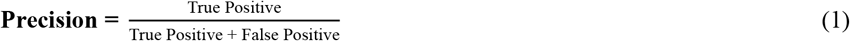

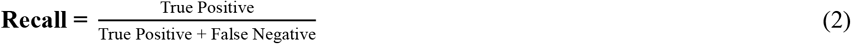

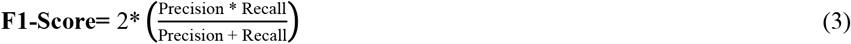

## 3. Results and Discussion

### 3.1 Performance Evaluation

To evaluate the performance of ANOMALY, we generated 50 simulated datasets containing 542 unique NuMTs using CHM13 human reference genome^13^. Our pipeline successfully detected a total 536 NuMTs (TP) across all samples, achieving an overall precision of 1, a recall of 0.989 and an F1-score of 0.994. The six NuMTs (FN) missed by the pipeline had size varied between 32 bp to 901 bp, with more than 66% NuMTs less than 100 bp (Supplementary Table 2, Supplementary Figure 1).

### 3.2 Resolution of Complex NuMT present in real-world data

To assess the pipeline’s performance on real-world data, we analysed long-read sequencing data from the NA12878 cell line (HG001)^22^ and compared the resulting NuMT calls with those derived from short-read data. ANOMALY identified 13 NuMTs, while DINUMT detected 11, with 8 calls shared between the two tools. Notably, ANOMALY failed to detect one true-positive NuMT identified by DINUMT, whereas DINUMT missed four true-positive calls and produced two false positives (Supplementary Table 3, Supplementary Figure 2).

ANOMALY also identified a discrepancy in interpreting the same NuMT event, whose nuclear breakpoint is located at chromosome 5:32,338,476. According to the short-read data, this NuMT measured 2,254 bp (MT:12723-14,977), whereas the long-read data revealed it to be only 291 bp in length (Supplementary Figure 3). Closer inspection showed that the NuMT insertion sequence maps to two distinct mitochondrial genome regions separated by 1,965 bp (MT:12723-12,867 and MT:14,832-14,977) (Supplementary Table 3). Because short-read-based NuMT callers rely on discordant read mapping, some discordant reads appeared to align with the first mitochondrial region. In contrast, others aligned to the second, resulting in an overestimation of the NuMT length. A schematic representation of this phenomenon is provided in Supplementary Figure 4. These findings highlight the importance of our tool that enables users to employ long-read sequencing data for accurately identifying complex NuMTs that the inherent length limitations of short-read sequencing data can misrepresent.

## 4. Conclusion

ANOMALY is a Snakemake pipeline for accurately identifying NuMTs from long-read sequencing data. It is an easy-to-install end-to-end pipeline that does not require an extensive experience with bioinformatics. In this article, we have displayed the use case of this pipeline on human data; however, this pipeline can be adapted to any organism as long as the long-read data is available. This pipeline also provides a publication-ready plot visualising NuMTs in the samples, which can be directly adapted to the research article.

## Acknowledgements

We thank the Functional Genomics Lab (DBEB, IIT Delhi) for proofreading and editing the manuscript.

## Availability and requirements

Project Name: ANOMALY.

Project Home Page: https://github.com/Nirmal2310/ANOMALY

Operating system(s): Linux.

Programming Language: Snakemake (Python), Bash, R.

Other requirements: Dependencies installed via conda and pip.

Licence: GNU GPLv3.

## References

1. Leister, D. Origin, evolution and genetic effects of nuclear insertions of organelle DNA. Trends Genet. 21, 655–663 (2005).

2. Wei, W. et al. Nuclear-embedded mitochondrial DNA sequences in 66,083 human genomes. Nature 611, 105–114 (2022).

3. Xue, L., Moreira, J. D., Smith, K. K. & Fetterman, J. L. The mighty NUMT: Mitochondrial DNA flexing its code in the nuclear genome. Biomolecules 13, (2023).

4. Turner, C. et al. Human genetic disease caused by de novo mitochondrial-nuclear DNA transfer. Hum. Genet. 112, 303–309 (2003).

5. Goldin, E. et al. Transfer of a mitochondrial DNA fragment to MCOLN1 causes an inherited case of mucolipidosis IV. Hum. Mutat. 24, 460–465 (2004).

6. Puertas, M. J. & González-Sánchez, M. Insertions of mitochondrial DNA into the nucleus-effects and role in cell evolution. Genome 63, 365–374 (2020).

7. Singh, K. K., Choudhury, A. R. & Tiwari, H. K. Numtogenesis as a mechanism for development of cancer. Semin. Cancer Biol. 47, 101–109 (2017).

8. Bensasson, D., Zhang, D.-X., Hartl, D. L. & Hewitt, G. M. Mitochondrial pseudogenes: evolution’s misplaced witnesses. Trends Ecol. Evol. 16, 314–321 (2001).

9. Dayama, G., Emery, S. B., Kidd, J. M. & Mills, R. E. The genomic landscape of polymorphic human nuclear mitochondrial insertions. Nucleic Acids Res. 42, 12640– 12649 (2014).

10. Wei, W. et al. Nuclear-mitochondrial DNA segments resemble paternally inherited mitochondrial DNA in humans. Nat. Commun. 11, 1740 (2020).

11. Kosugi, S. & Terao, C. Comparative evaluation of SNVs, indels, and structural variations detected with short- and long-read sequencing data. Hum. Genome Var. 11, 18 (2024).

12. Köster, J. & Rahmann, S. Snakemake--a scalable bioinformatics workflow engine. Bioinformatics 28, 2520–2522 (2012).

13. Nurk, S. et al. The complete sequence of a human genome. Science 376, 44–53 (2022).

14. Smolka, M. et al. Detection of mosaic and population-level structural variants with Sniffles2. Nat. Biotechnol. 42, 1571–1580 (2024).

15. Ye, J., McGinnis, S. & Madden, T. L. BLAST: improvements for better sequence analysis. Nucleic Acids Res. 34, W6–9 (2006).

16. Shen, W., Le, S., Li, Y. & Hu, F. SeqKit: A cross-platform and ultrafast toolkit for FASTA/Q file manipulation. PLoS One 11, e0163962 (2016).

17. Li, H. et al. The Sequence Alignment/Map format and SAMtools. Bioinformatics 25, 2078–2079 (2009).

18. Quinlan, A. R. & Hall, I. M. BEDTools: a flexible suite of utilities for comparing genomic features. Bioinformatics 26, 841–842 (2010).

19. Logsdon, G. A., Vollger, M. R. & Eichler, E. E. Long-read human genome sequencing and its applications. Nat. Rev. Genet. 21, 597–614 (2020).

20. Gu, Z., Gu, L., Eils, R., Schlesner, M. & Brors, B. circlize Implements and enhances circular visualisation in R. Bioinformatics 30, 2811–2812 (2014).

21. Ono, Y., Hamada, M. & Asai, K. PBSIM3: a simulator for all types of PacBio and ONT long reads. NAR Genom. Bioinform. 4, lqac092 (2022).

22. Jain, M. et al. Nanopore sequencing and assembly of a human genome with ultra-long reads. Nat. Biotechnol. 36, 338–345 (2018).

