## Supplementary Figures for "ANOMALY: A Snakemake pipeline for identifying NuMTs from Long-Read Sequencing Data"

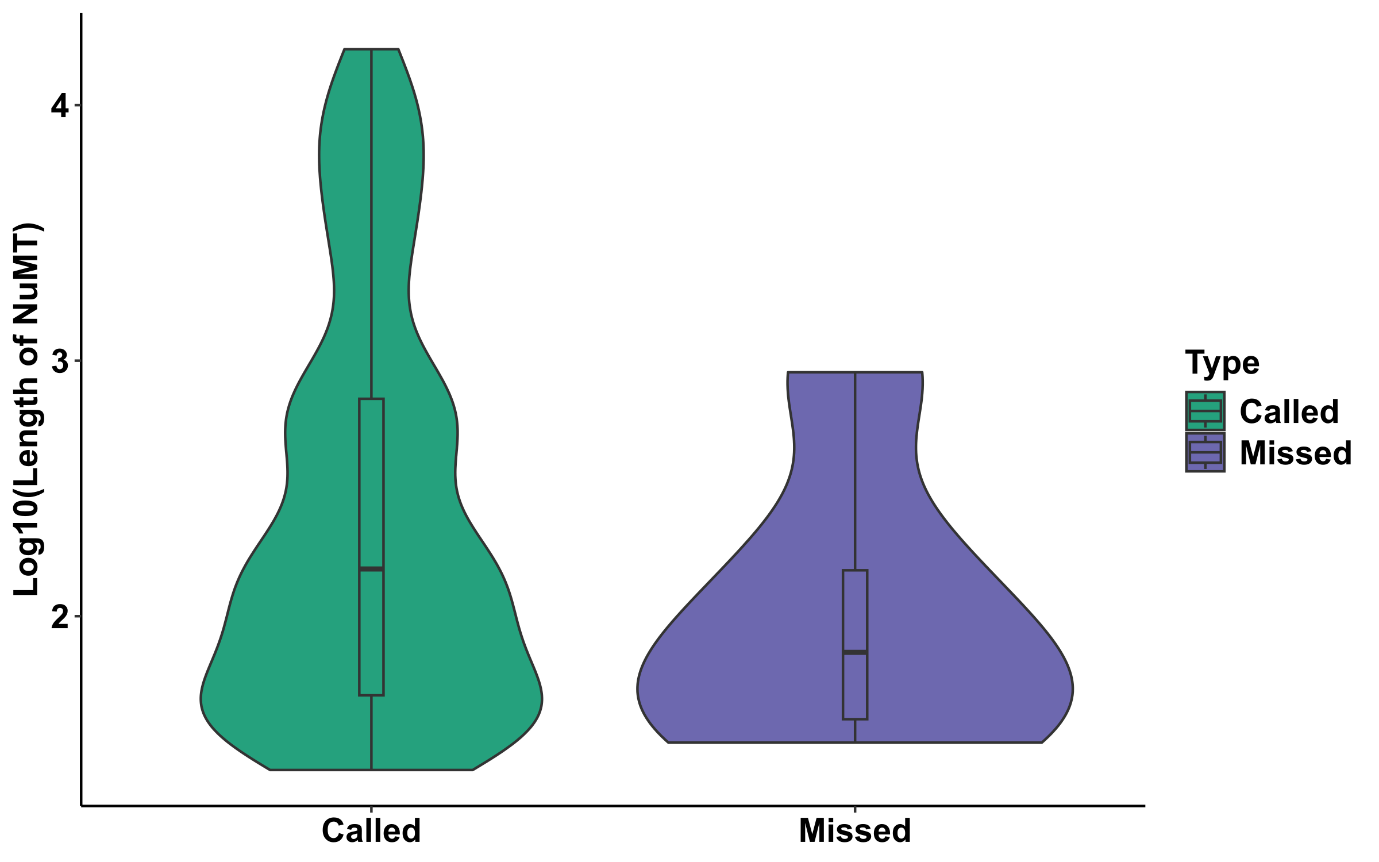

**Supplementary Figure 1:** Violin Plot showing the length distribution of NuMTs called and missed by the pipeline. 4 out of 6 missed NuMTs had a length of less than 100 bp.

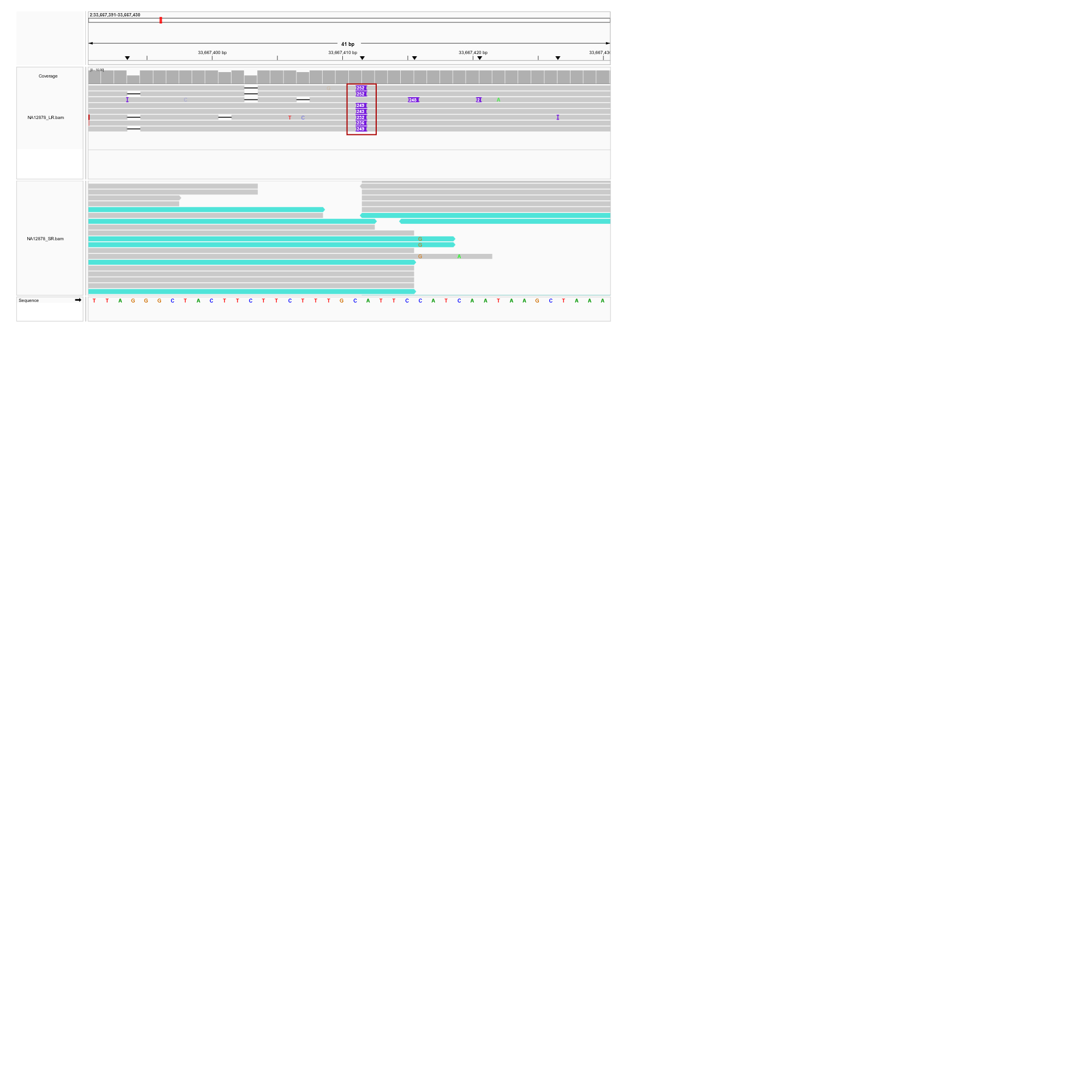

**Supplementary Figure 2(A):** IGV Screenshot of an NuMT called by both ANOMALY and DINUMT. The NuMT is shown as an insertion in Long-read sequencing data and as discordant reads mapping to mitochondrial genome (turquoise colour) in Short-read sequencing data.

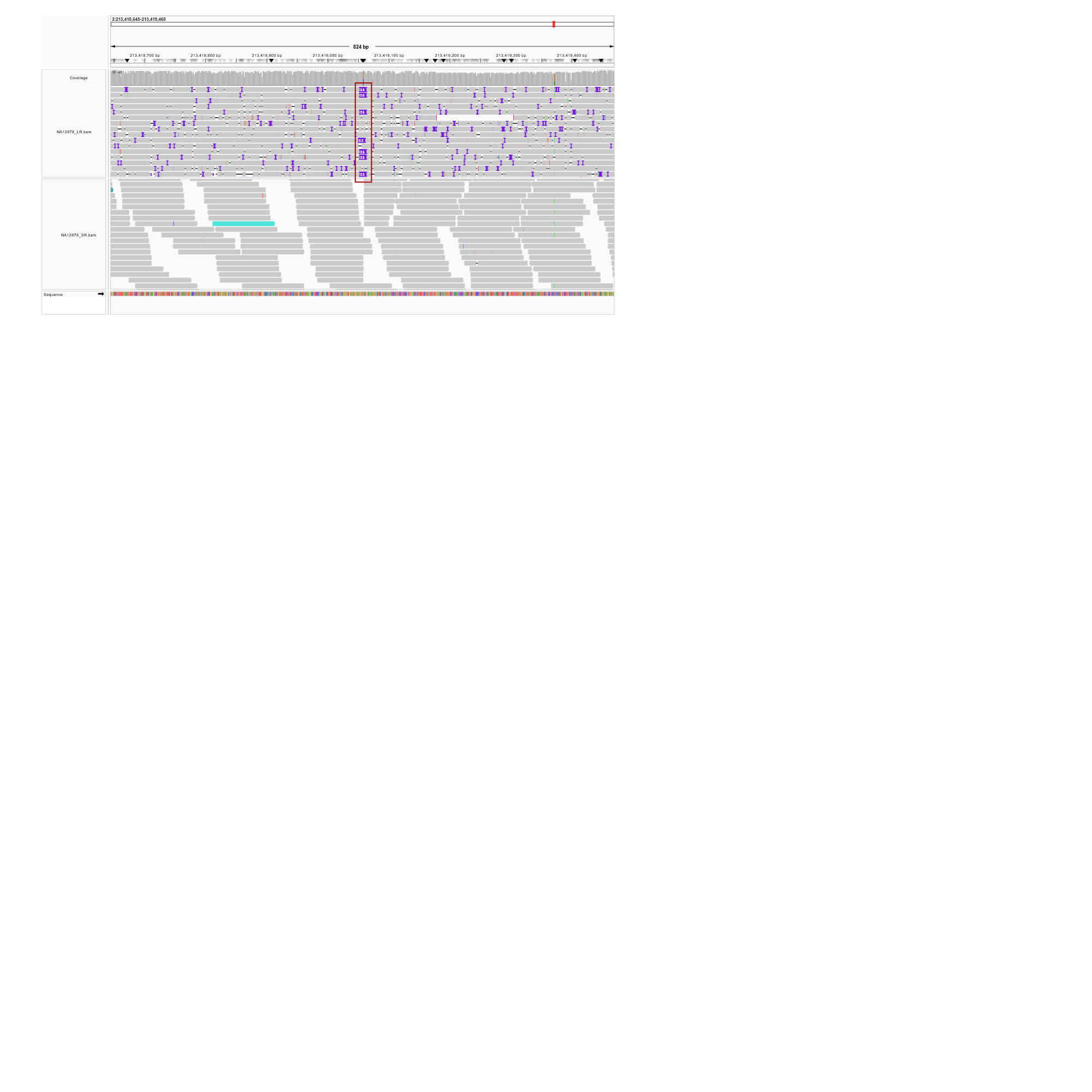

**Supplementary Figure 2(B):** IGV Screenshot of an NuMT called by both ANOMALY and DINUMT. The NuMT is shown as an insertion in Long-read sequencing data and as discordant reads mapping to mitochondrial genome (turquoise colour) in Short-read sequencing data.

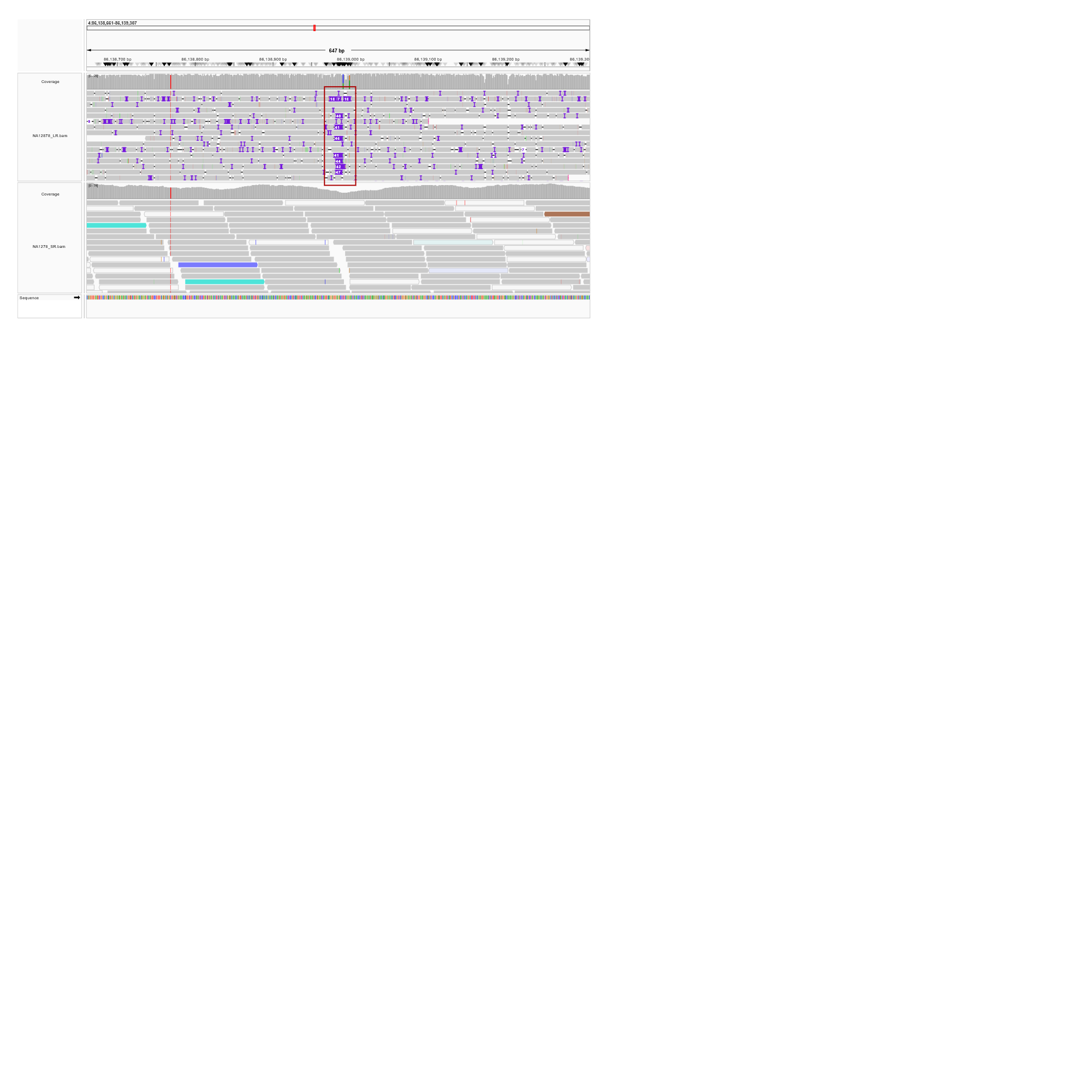

**Supplementary Figure 2(C):** IGV Screenshot of an NuMT called by ANOMALY only. The NuMT is shown as an insertion in Long-read sequencing data and as discordant reads mapping to mitochondrial genome (turquoise colour) in Short-read sequencing data. Even after the presence of discordant reads, this NuMT call is missed by DINUMT.

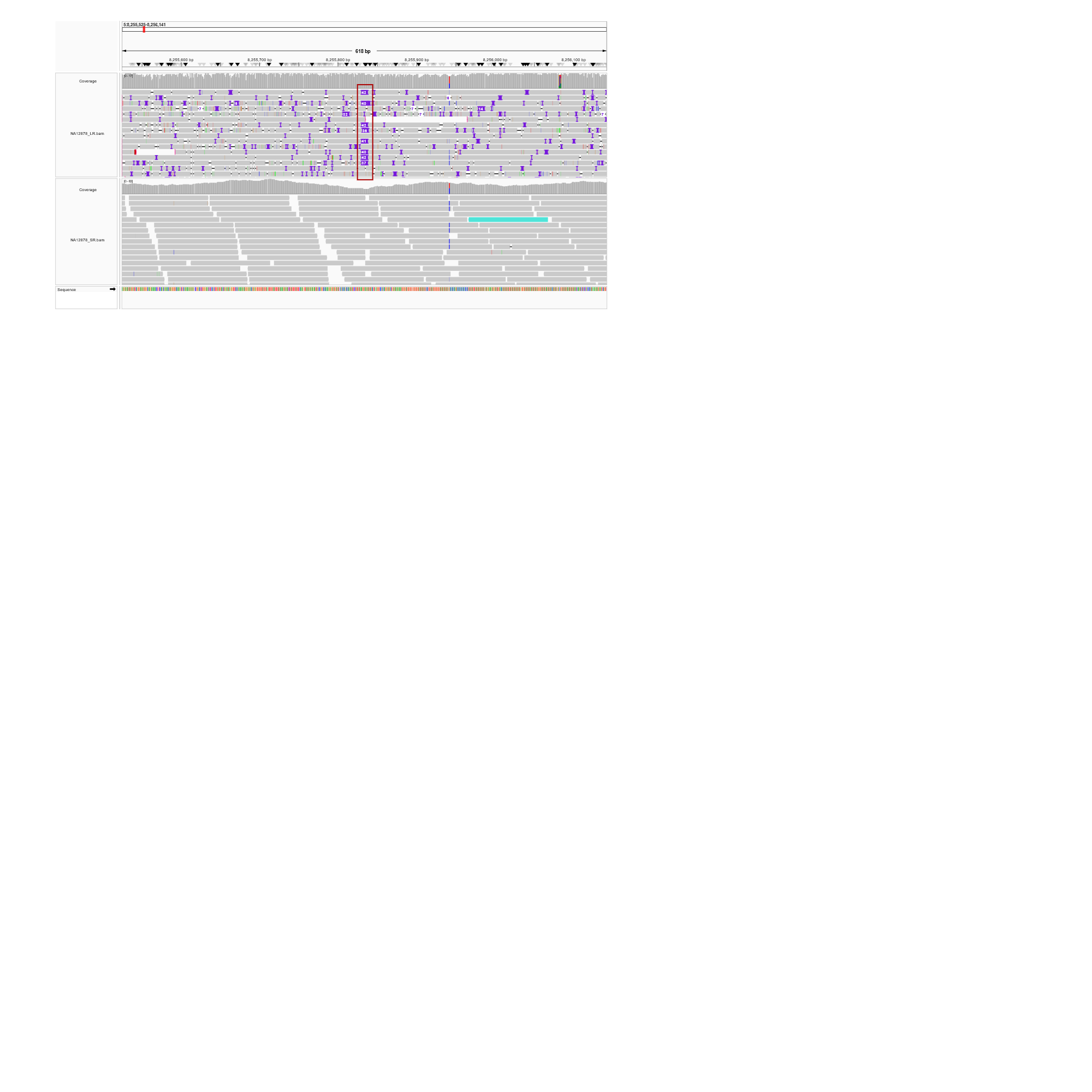

**Supplementary Figure 2(D):** IGV Screenshot of an NuMT called by both ANOMALY and DINUMT. The NuMT is shown as an insertion in Long-read sequencing data and as discordant reads mapping to mitochondrial genome (turquoise colour) in Short-read sequencing data.

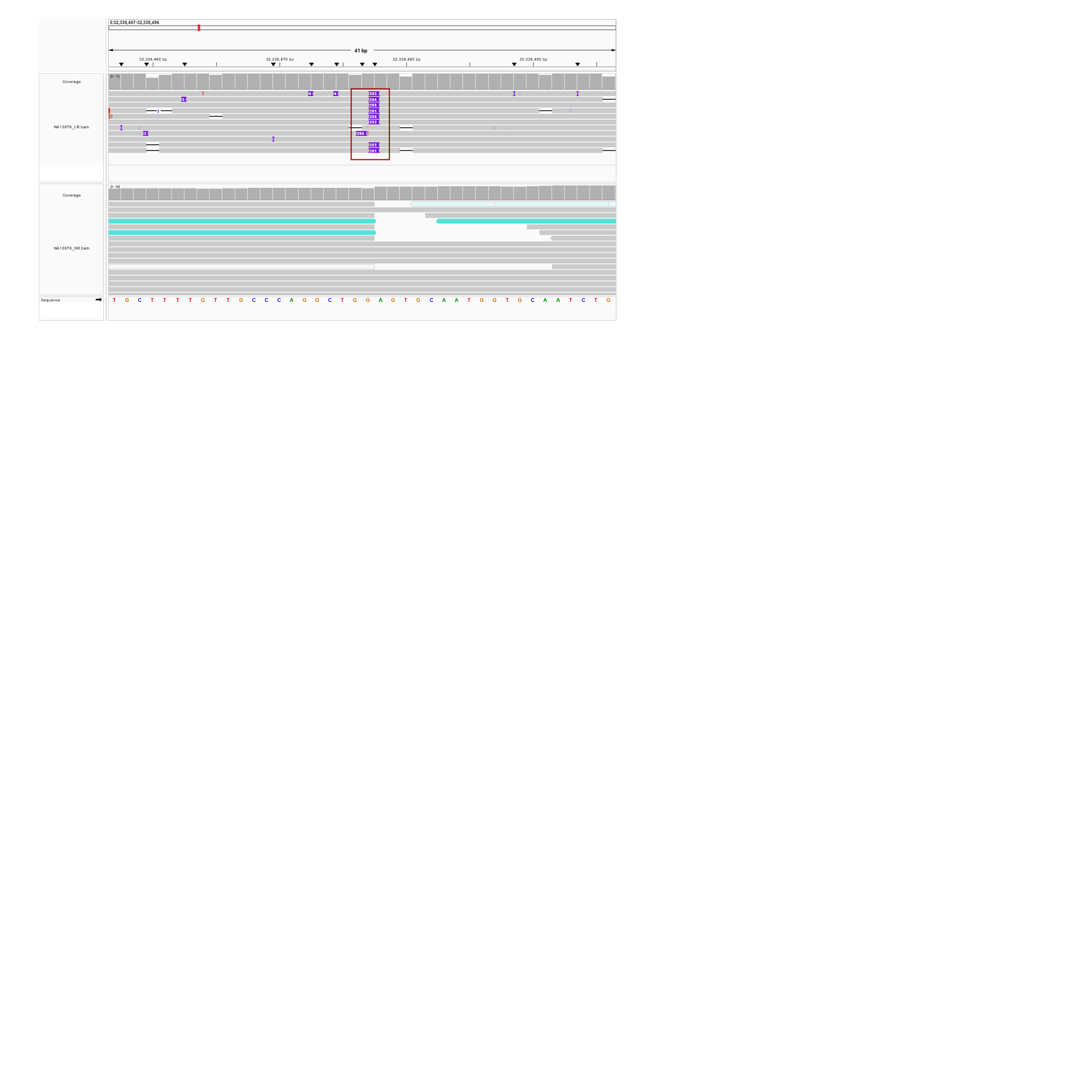

**Supplementary Figure 2(E):** IGV Screenshot of an NuMT called by both ANOMALY and DINUMT. The NuMT is shown as an insertion in Long-read sequencing data and as discordant reads mapping to mitochondrial genome (turquoise colour) in Short-read sequencing data.

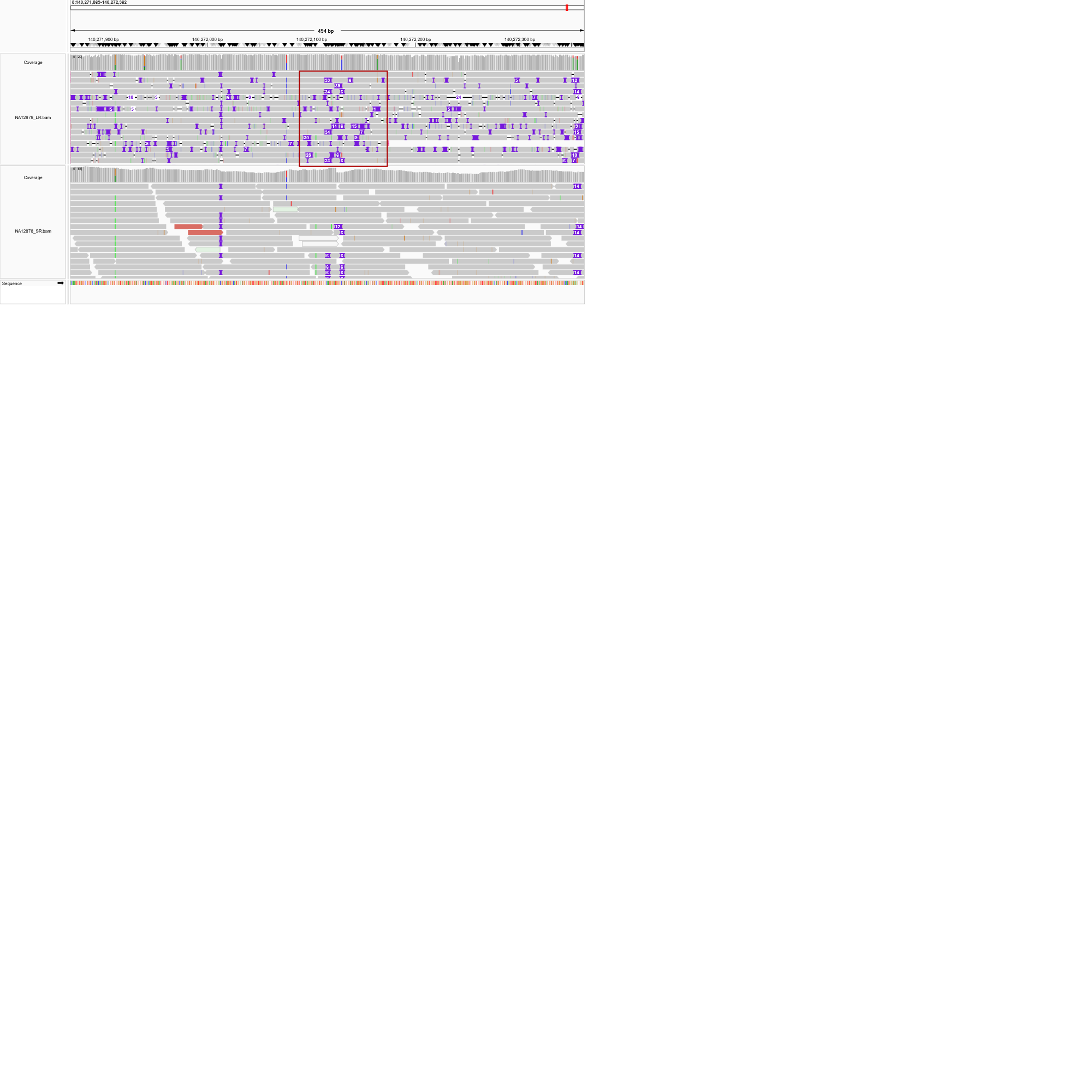

**Supplementary Figure 2(F):** IGV Screenshot of an NuMT called by ANOMALY only. The NuMT is shown as an insertion in Long-read sequencing data. This NuMT is completely missed by DINUMT.

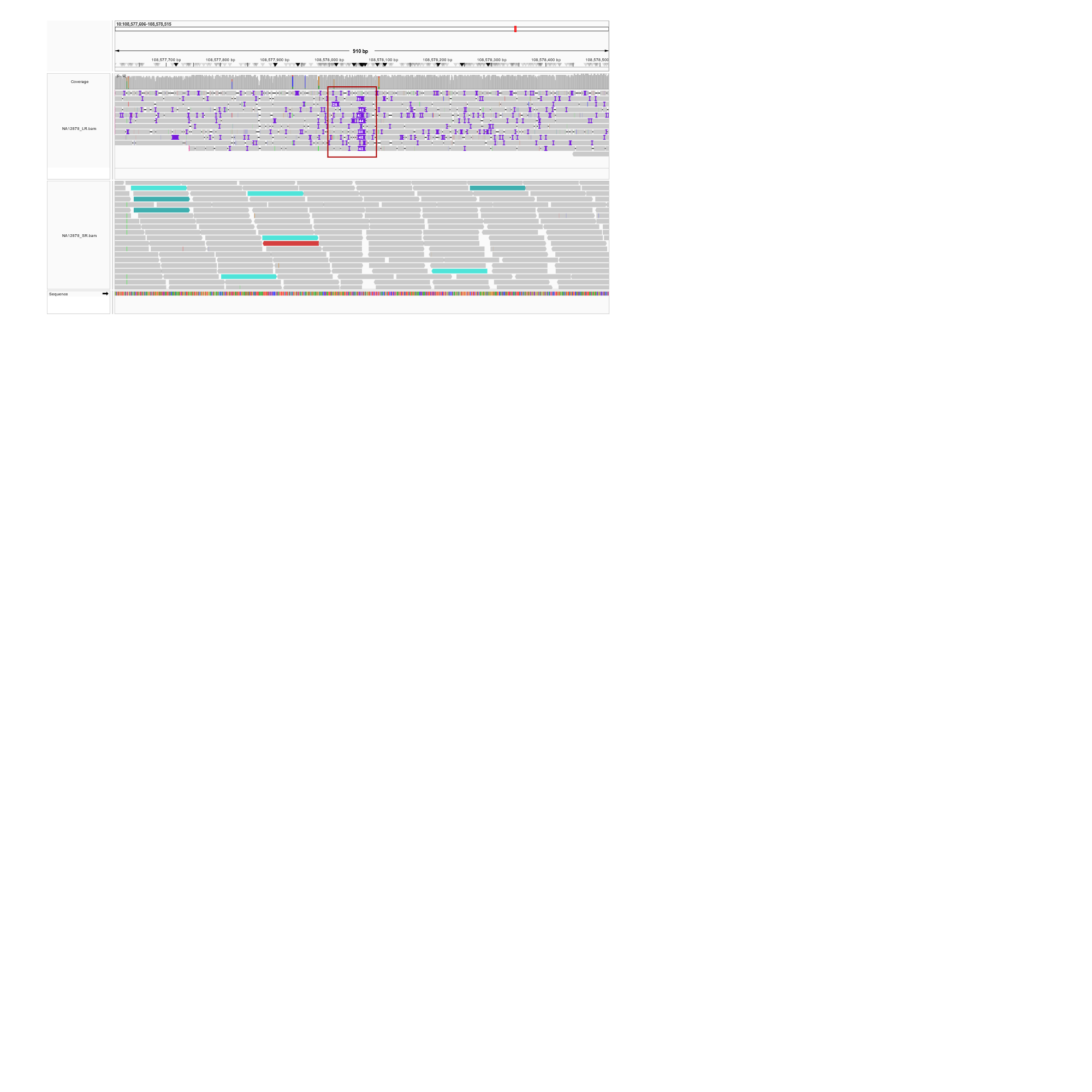

**Supplementary Figure 2(G):** IGV Screenshot of an NuMT called by both ANOMALY and DINUMT. The NuMT is shown as an insertion in Long-read sequencing data and as discordant reads mapping to mitochondrial genome (turquoise colour) in Short-read sequencing data.

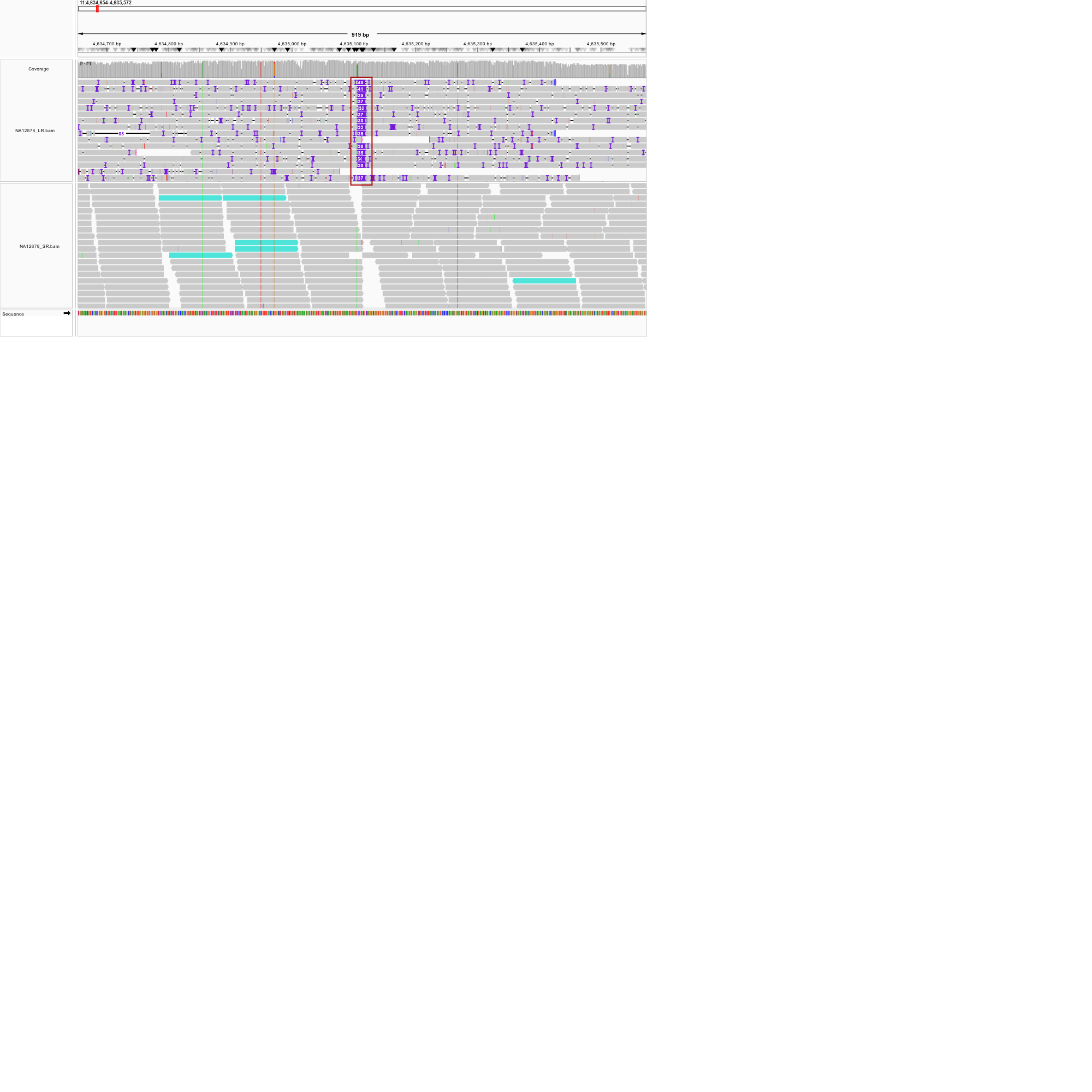

**Supplementary Figure 2(H):** IGV Screenshot of an NuMT called by both ANOMALY and DINUMT. The NuMT is shown as an insertion in Long-read sequencing data and as discordant reads mapping to mitochondrial genome (turquoise colour) in Short-read sequencing data.

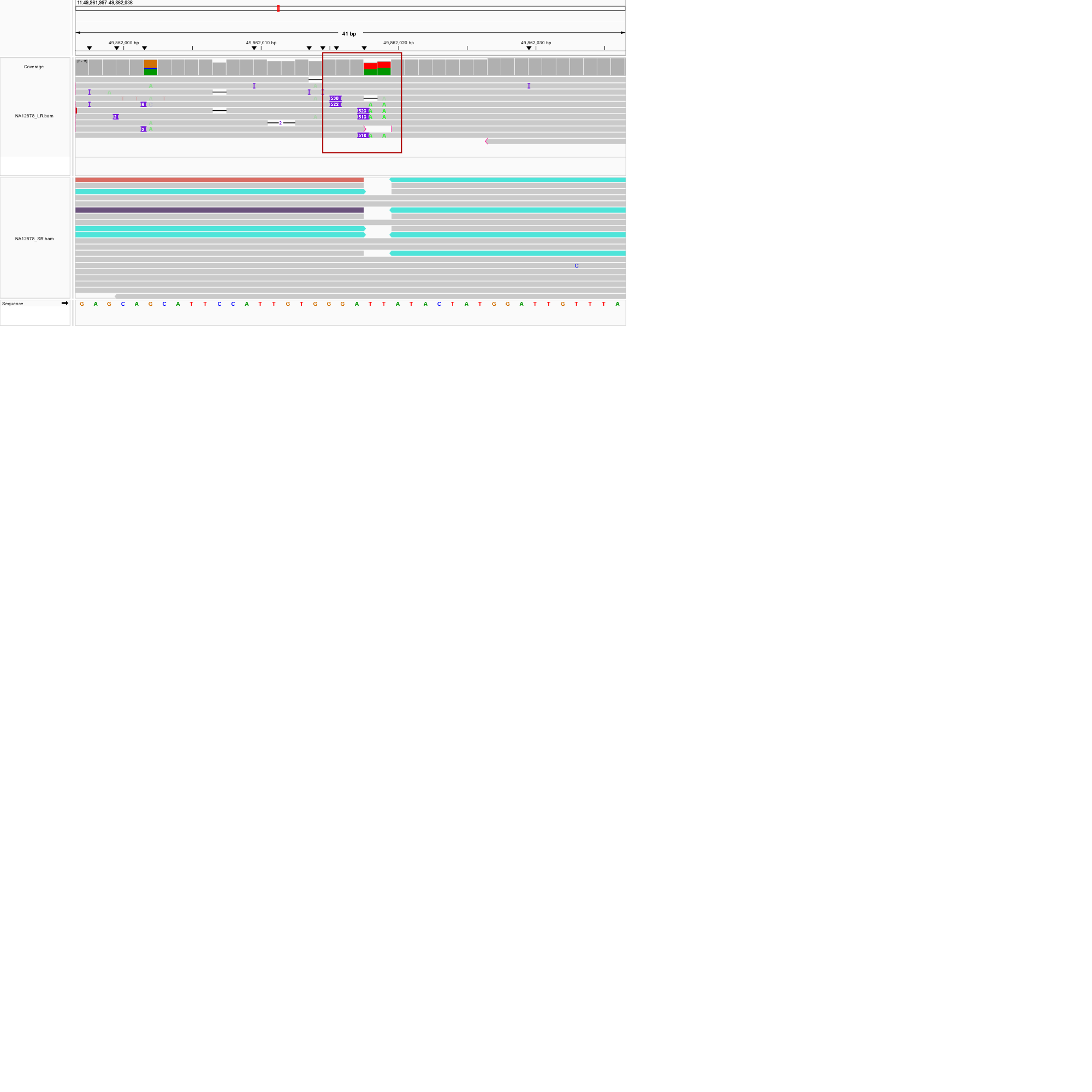

**Supplementary Figure 2(I):** IGV Screenshot of an NuMT called by both ANOMALY and DINUMT. The NuMT is shown as an insertion in Long-read sequencing data and as discordant reads mapping to mitochondrial genome (turquoise colour) in Short-read sequencing data.

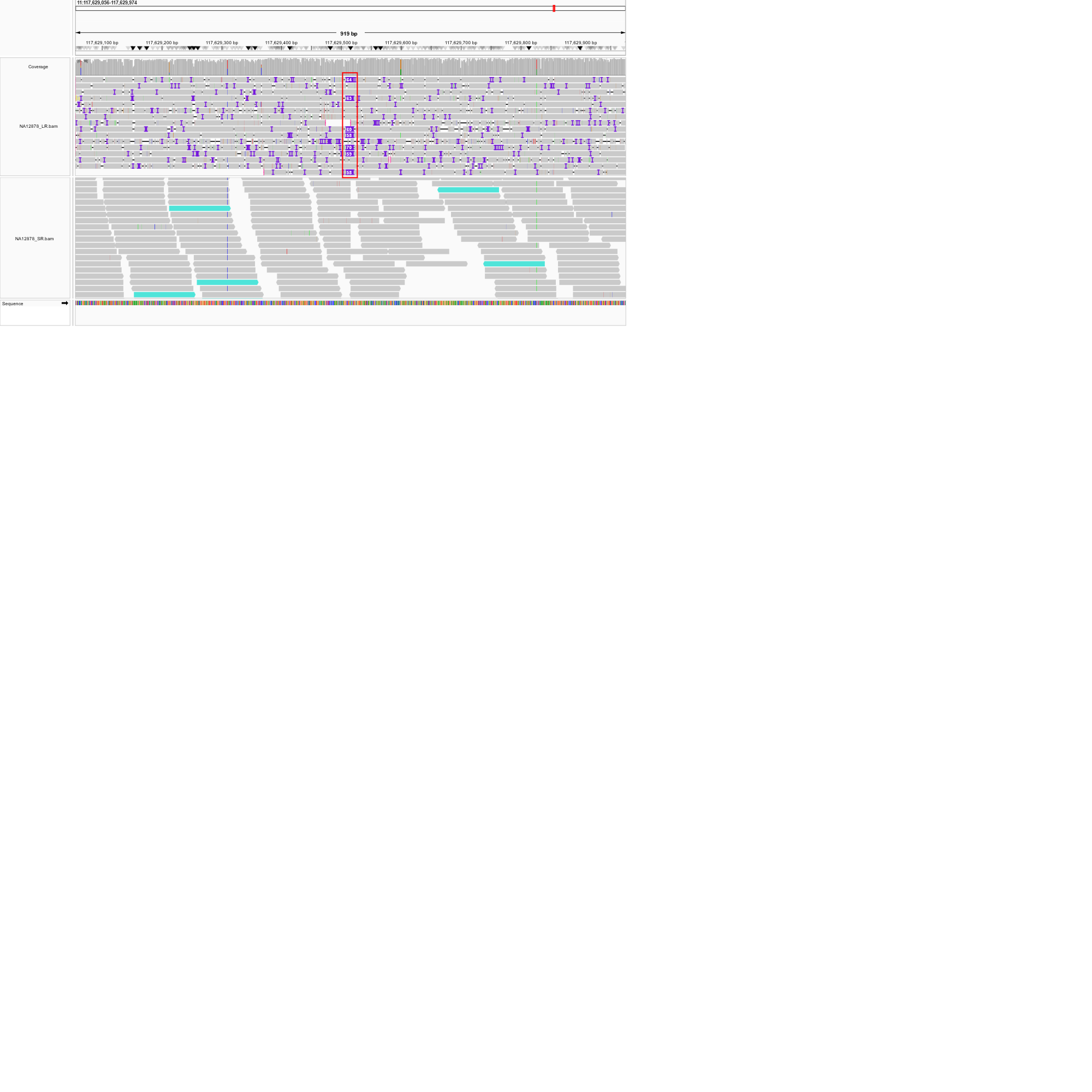

**Supplementary Figure 2(J):** IGV Screenshot of an NuMT called by both ANOMALY and DINUMT. The NuMT is shown as an insertion in Long-read sequencing data and as discordant reads mapping to mitochondrial genome (turquoise colour) in Short-read sequencing data.

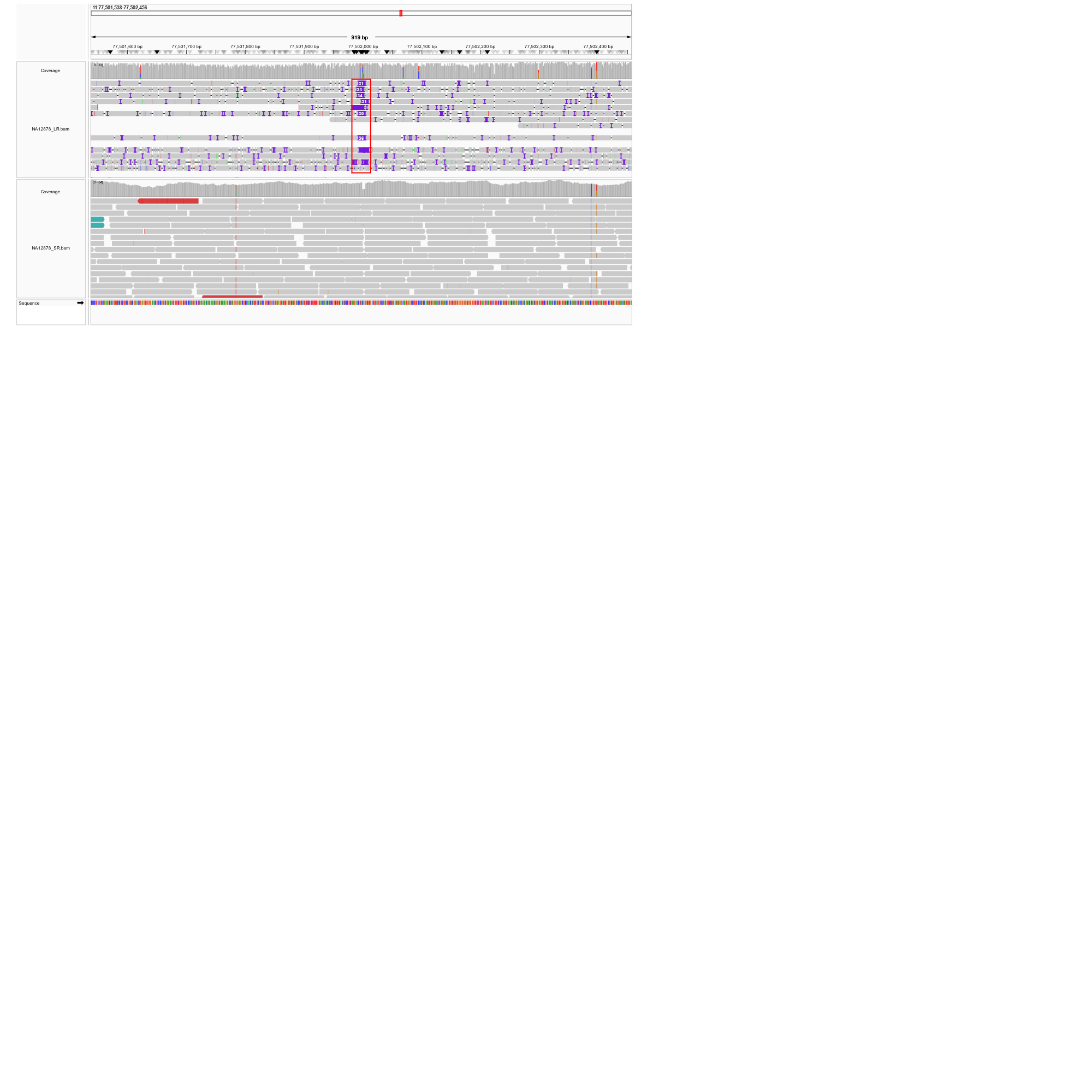

**Supplementary Figure 2(K):** IGV Screenshot of an NuMT called by DINUMT and missed by ANOMALY. The NuMT is shown as an insertion in Long-read sequencing data but the pipeline missed this call because of the high E-value.

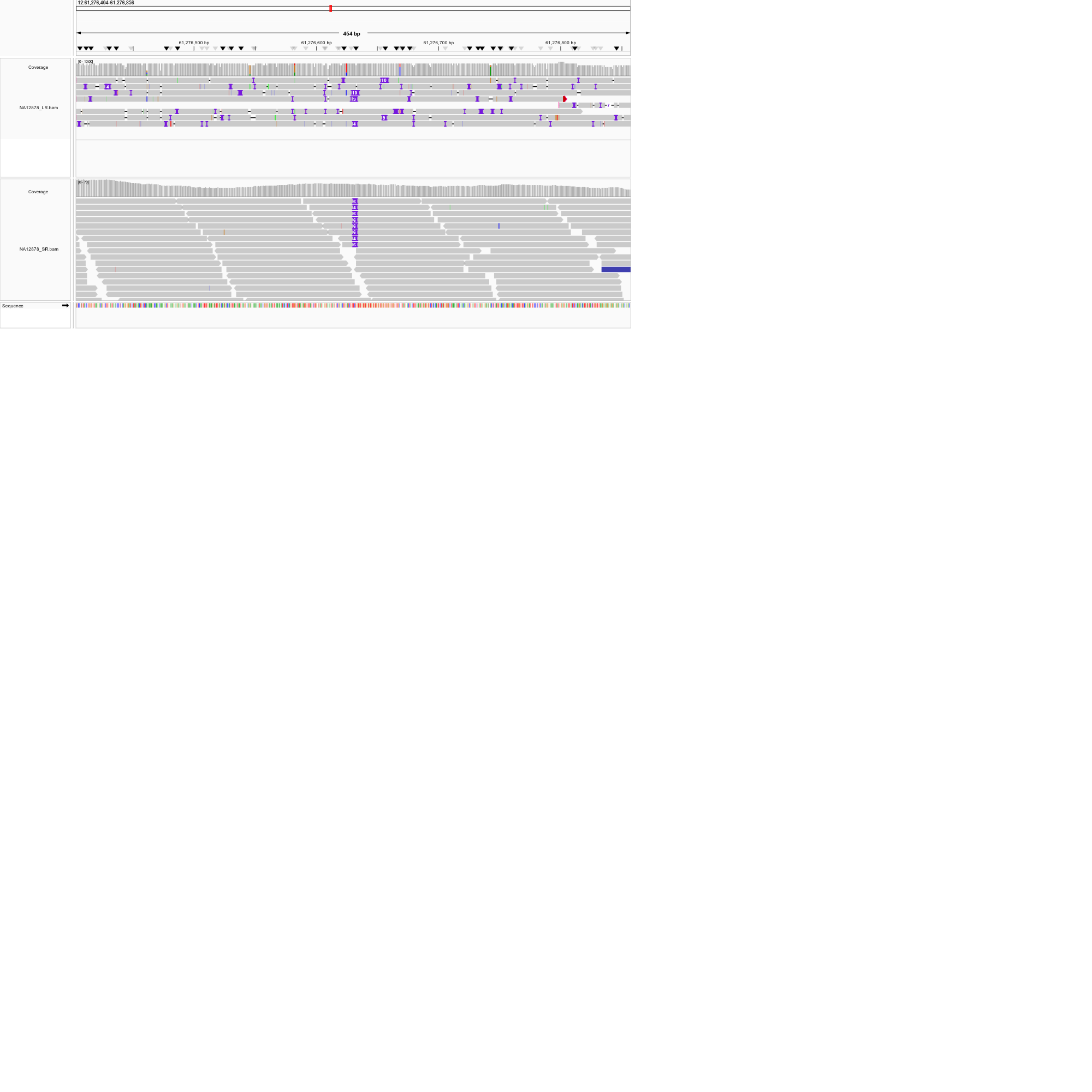

**Supplementary Figure 2(L):** IGV Screenshot of an NuMT called by DINUMT only. This call is a False-Positive as there are no signatures of insertions in both long-read as well as short-read sequencing data.

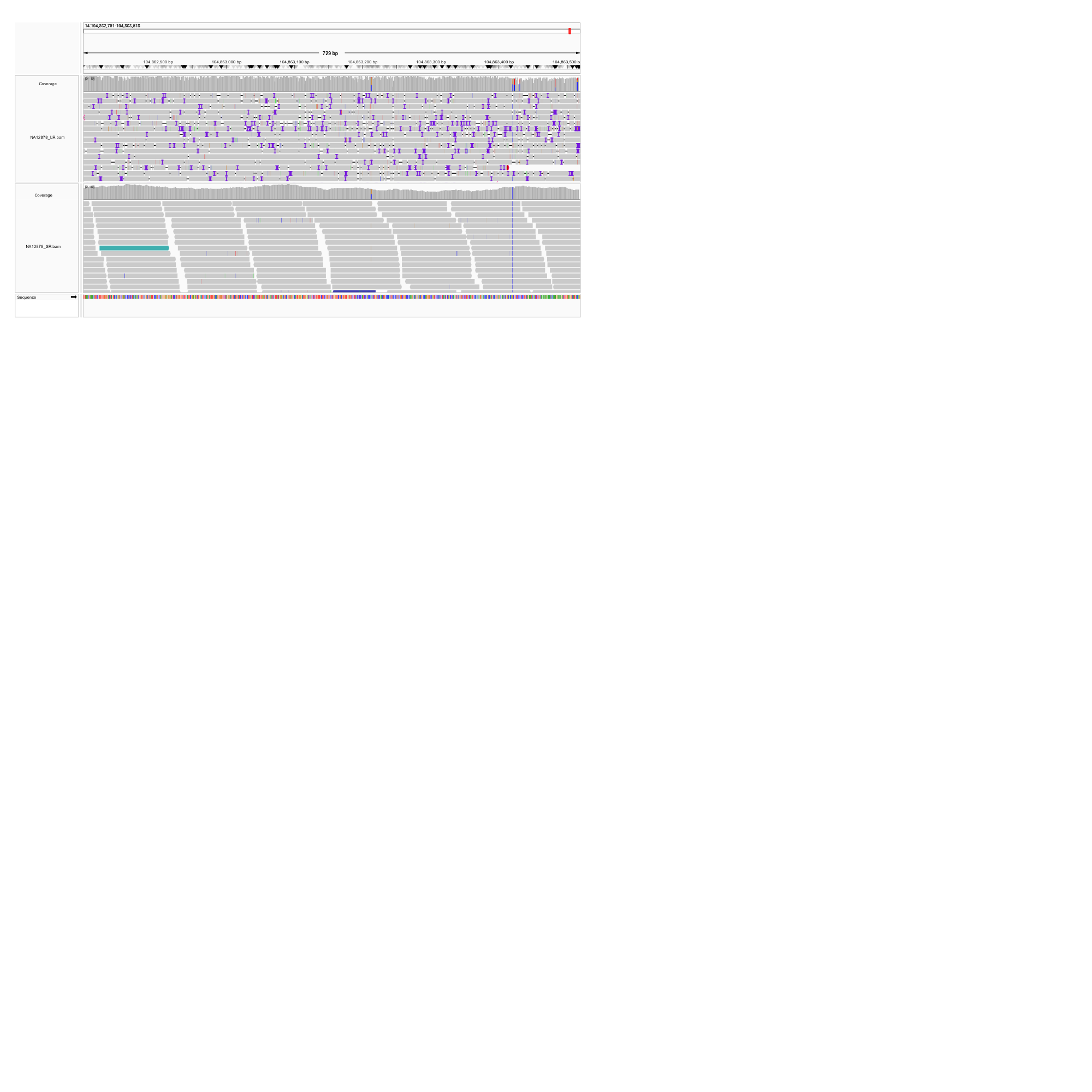

**Supplementary Figure 2(M):** IGV Screenshot of an NuMT called by DINUMT only. This call is a False-Positive as there are no signatures of insertions in both long-read as well as short-read sequencing data.

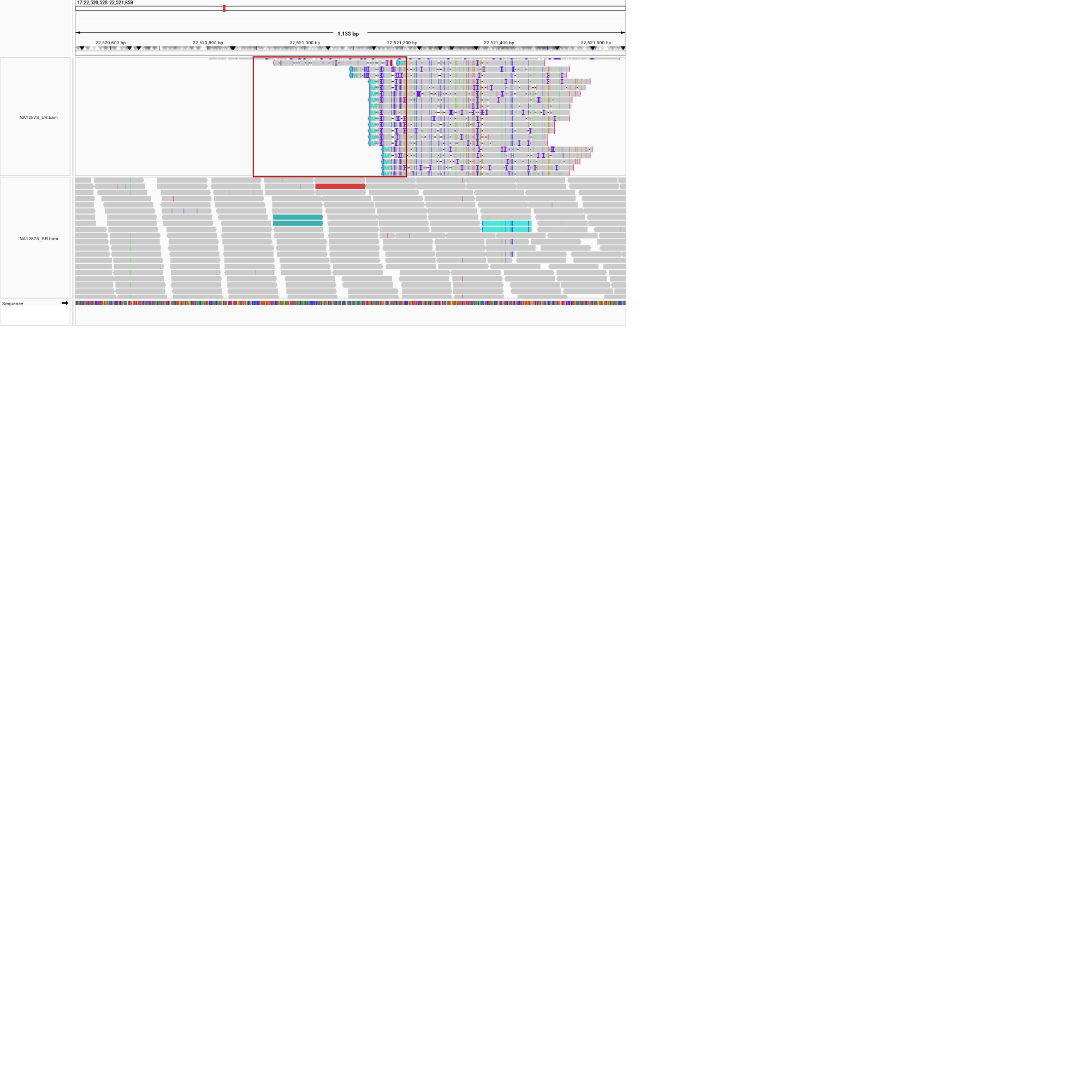

**Supplementary Figure 2(N):** IGV Screenshot of an NuMT called by ANOMALY only. The NuMT is shown as soft-clipped reads Long-read sequencing data. This NuMT is completely missed by DINUMT even though discordant reads mapped to mitochondrial genome are present in short-read sequencing data.

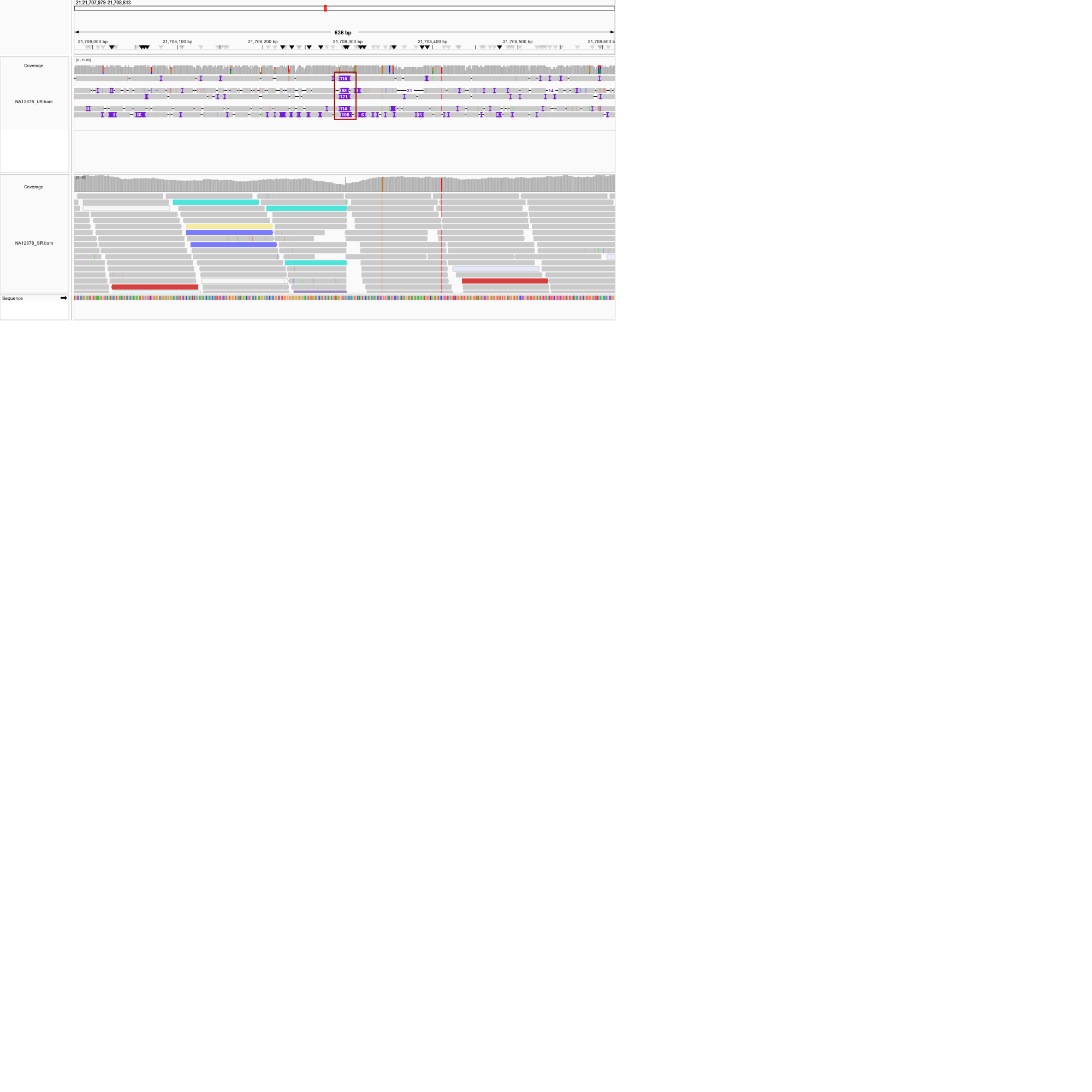

**Supplementary Figure 2(O):** IGV Screenshot of an NuMT called by ANOMALY only. The NuMT is shown as an insertion in Long-read sequencing data. This NuMT is completely missed by DINUMT even though discordant reads mapped to mitochondrial genome are present in short-read sequencing data.

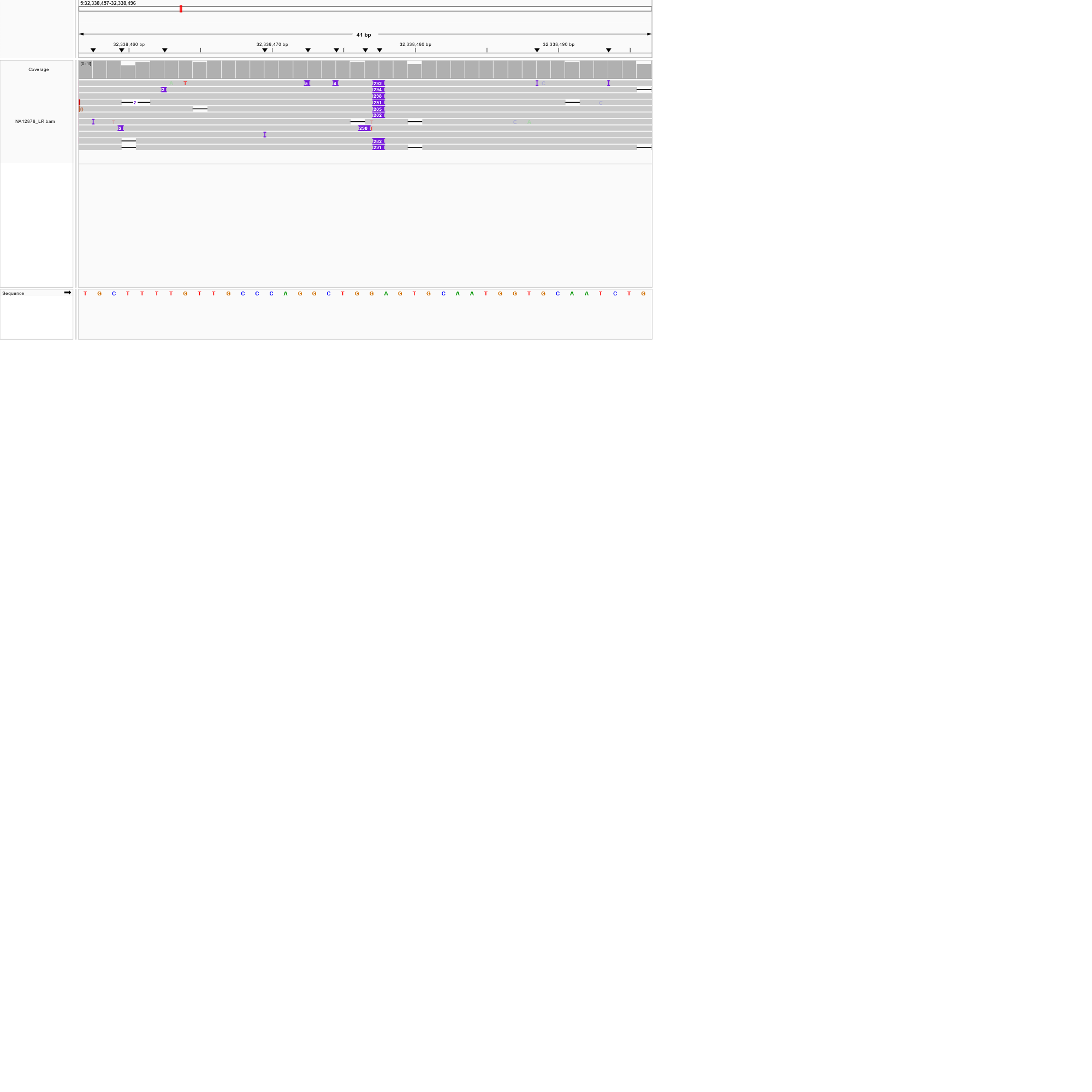

**Supplementary Figure 3:** IGV Screenshot showing the NuMT insertion in the long-read sequencing data of the NA12878 cell line. The NuMT shows a discrepancy when called using short-read data, as its size increases from 291 bp to 2,254 bp.

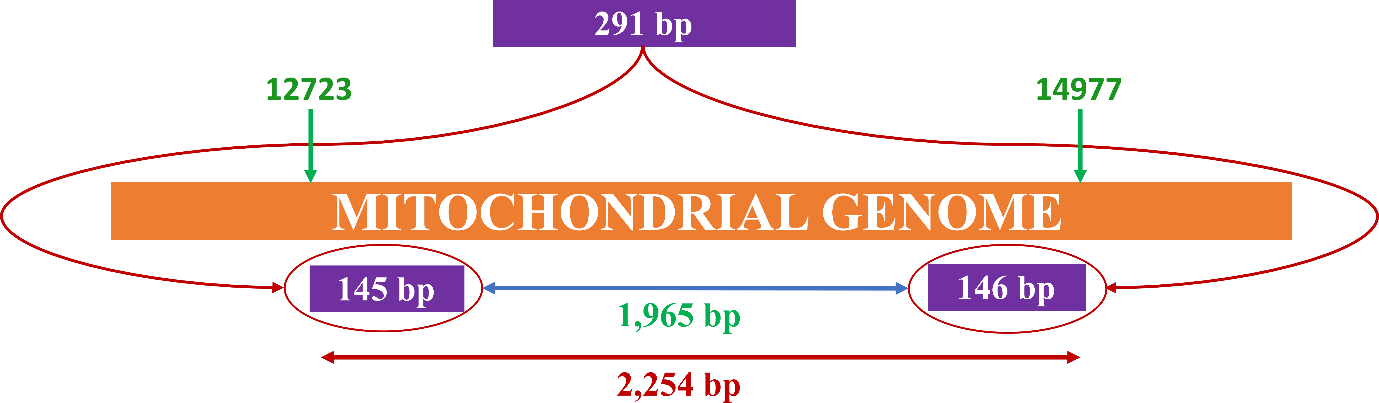

**Supplementary Figure 4:** Schematic Representation of a unique NuMT found in HG001 (NA12878) cell line that shows discrepancy when called using short-read sequencing data and long-read sequencing data. The short-read-based caller identified the NuMT with the size of 2,254 bp; however, the actual size is 291 bp, as identified using the long-read data. This might be because the NuMT is mapping to two parts of the mitochondrial genome that are 1,965 bp apart. Since short-read-based callers utilise discordant reads to identify the NUMTs, some reads might be mapped to the first part of the mitochondrial genome, and others might be mapped to the second part. Discordant reads mapping to two parts of the genome might be the main cause of the short-read-based callers' overestimation of NuMT size.
